# Up sampled PSF enables high accuracy 3D super-resolution imaging with sparse sampling rate

**DOI:** 10.1101/2024.10.31.620776

**Authors:** Jianwei Chen, Wei Shi, Jianzheng Feng, Jianlin Wang, Sheng Liu, Yiming Li

## Abstract

Single-molecule localization microscopy (SMLM) provides nanoscale imaging, but pixel integration of acquired SMLM images limited the choice of sample rate which restricts the information content conveyed within each image. We propose an up-sampled point spread function (PSF) inverse modeling method for large-pixel single molecule localization, enabling precise 3D super-resolution imaging with sparse sampling rate. Our approach could reduce data volume or expand the field of view by nearly an order of magnitude, while maintaining high localization accuracy, greatly improving the imaging throughput with limited pixels offered by the existing cameras.

---

Single-molecule localization microscopy (SMLM) has emerged as one of the most powerful techniques for surpassing the diffraction limit of light microscopy, enabling the visualization of biological structures at the nanoscale^1,2^. However, achieving one super resolved SMLM image typically demands the acquisition of tens of thousands of single molecule images. This is particularly challenging in reproducible biological research, where statistically robust results require large datasets. The sheer volume of data generated not only requires significant computational power but also demands extensive storage management, creating heavy burden in both data processing and resource allocation especially when large number of SMLM images are needed^3–7^.

A fundamental aspect of SMLM involves the precise localization of diffraction-limited singe molecule spots, typically captured by cameras as pixelated images. Large-pixel imaging, which is simply achieved by lowering the microscope magnification, could largely reduce the data volume with the same field of view (FOV). It has long been noticed that pixelation effect becomes especially problematic when the pixel size exceeds the ideal sampling rate (∼100 nm) for single molecule localization under high numerical aperture (NA) objective^8–10^. Recent innovations like experimental spline-interpolated^11,12^ and in-situ PSF modeling^13^ have helped achieve the Cramér-Rao lower bound (CRLB) for single molecule localization. However, effective algorithms to fully addresses pixelation noise are still missing in the field, forcing researchers to compromise on pixel size, limiting image quality and data content.

Empirical observations suggest that varying the integration positions of bead data (Figure 2a-b) can retain continuous PSF information, which is otherwise lost in pixelated images. This discovery has led to the development of our inverse integration technique, which could estimate a pre-integrated, up-sampled PSF from pixelated data. By recovering high-frequency details that are normally blurred by pixelation, this up-sampled PSF significantly enhances localization precision for under-sampled data. As a result, our approach not only allows for optimal localization accuracy as defined by the CRLB, but also mitigates the computational and storage challenges associated with large-scale SMLM. By reducing the widely used sampling rate for high NA objective by 3 times, we could decrease the data volume by 89% or expanding the field of view by nine-fold. Our method facilitates more efficient high-precision imaging which normally requires fine sampling, addressing the pressing need for computationally accessible solutions in reproducible large-scale biological studies with super resolution.

The up sampled PSF inverse modeling process involves reconstructing the up sampled PSF from pixelated beads data, a common inverse problem. In microscopy, the intensity field from bead fluorescence is integrated on the camera chip to form an image, making forward modeling straightforward when parameters such as the emission position and the up sampled 3D PSF are known. However, the inverse process of determining the up sampled PSF is much more complex, requiring sufficient sampling of different bead positions and an optimization algorithm to retrieve the underlying shared continuous PSF model. Our up sampled PSF modeling process (Figure 1) starts by extracting regions of interest (ROIs) from experimental beads z-stack data, generating a set of pixelated PSFs randomly distributed at different positions. An initial up sampled PSF model, along with parameters like 3D position, photon counts, and background, is used to generate fine PSFs. The fine PSFs are normally integer multiple times up sampled from the under sampled coarse PSFs. Therefore, we bin the fine PSFs over the pixel size to create coarse PSFs, which are fitted to the experimental data based on Poisson maximum likelihood estimation (MLE)^14^. The optimization algorithm L-BFGS-B^15^ is then employed to iteratively adjust the up sampled PSF model and related parameters (SI Note 1).

**Figure 1.**
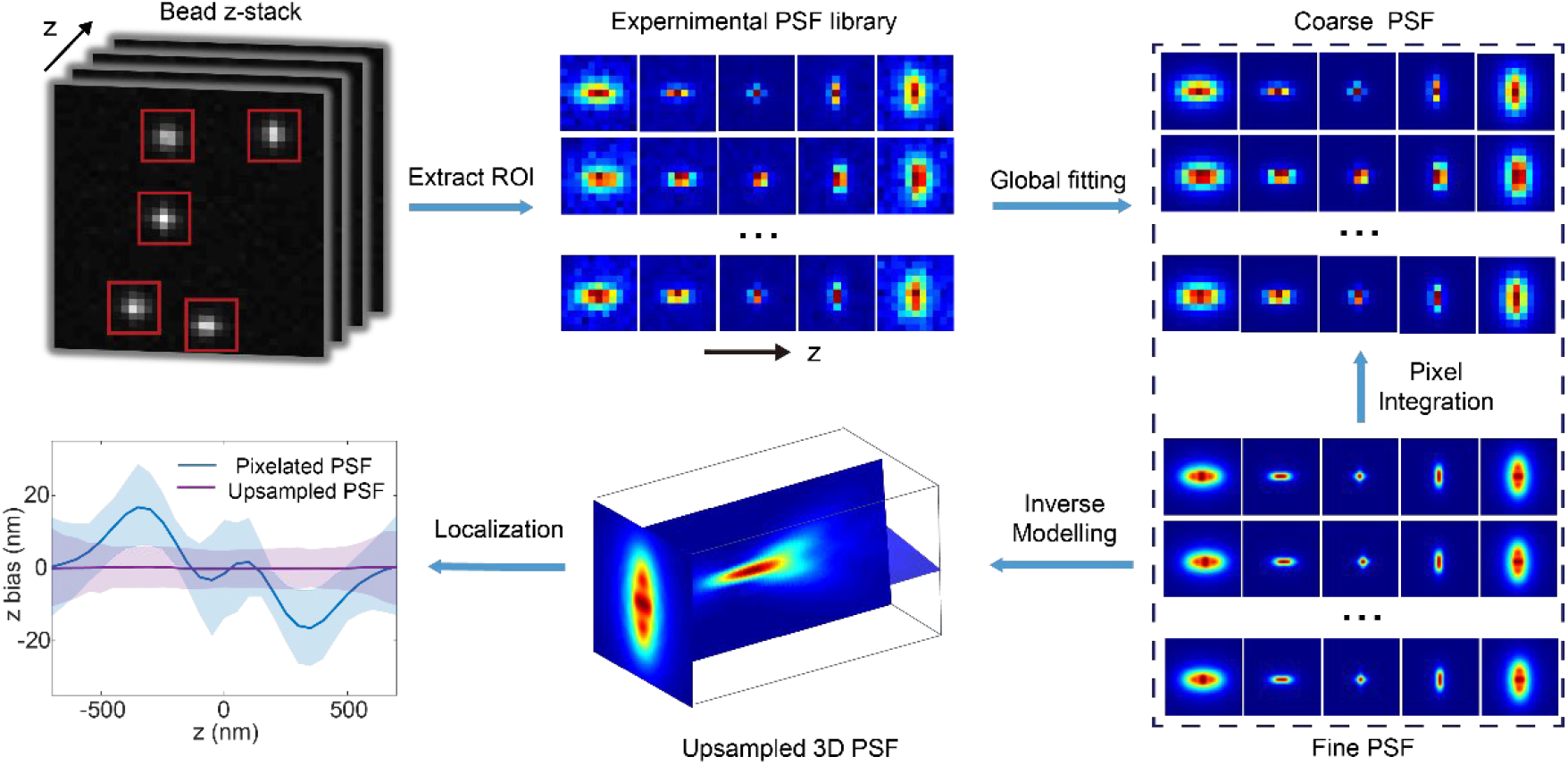
Concept of up sampled PSF modelling. Beads z-stack data (Top left) were first collected to extract ROIs and build the experimental PSF library locating at different positions (Top center). MLE was performed to the experimental beads data library to retrieve the global shared up sampled PSF. Here, the coarse PSF was obtained by pixel integration of the up sampled PSF. We finally localize the experimental bead data with both the un sampled PSF (Bottom center) and averaged coarse PSF, revealing that only the up sampled PSF could achieve unbiased and optimal localization results (Bottom left).

After obtaining the up sampled PSF from the beads data, we then developed a new localization algorithm which could minimize the localization bias for pixelated single molecule data with large pixel size. For up sampled PSF localization, spline interpolated PSF was employed so that it is applicable to PSFs with arbitrary shape. Different to the conventional spline PSF fitting, we introduced a new input constant, “bin,” to perform pixel integration, allowing for precise localization across different sampling rates between the up sampled PSF and the single molecule data (SI Note 2). Prior to localization, the up sampled PSF is preprocessed by convolving it with a *bin × bin ker*nel with all elements as 1 (ED Fig 1a, SI Note 2). This step is essential since the experimental data and the up sampled PSF are integrated over different pixel sizes, leading to shape discrepancies. After convolution-based preprocessing, spline interpolation (SI Note 2) is applied to the up sampled PSF and used for bead or single-molecule localization. Compared with the conventional spline fitter without considering the pixel integration effect, we showed that our up sampled spline fitting algorithm could achieve CRLB at different pixel sizes and effectively reduced the localization bias by a factor of 2 for 330 nm pixel size data (ED Fig 2).

We then systematically investigated the optimal pixel size for large-pixel imaging by calculating the theoretical CRLB for PSFs of varying pixel sizes using vectorial PSF model^16^ (SI Note 3 and 4). As shown in ED Fig 3, CRLB of large pixel PSF is spatially variant. To accurately reflect the localization precision of large-pixel PSF, we uniformly selected 121 points within a single pixel, calculated the CRLB for each point, and averaged these values to obtain the final mean CRLB (ED Fig 4a-c). As shown in Figure 2c and ED Fig 4a-c, compared to the traditional 110 nm pixel size PSF, the 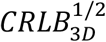^17^ of a 220 nm pixel size PSF increased by only about 6.7% for astigmatic PSF^18^, while the 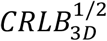 *for* a 330nm pixel size PSF increased by just 18.9%. Therefore, a factor of 2-3 times increase of traditional pixel size (∼ 100 nm) would greatly reduce the data volume while maintaining high localization precision.

**Figure 2.**
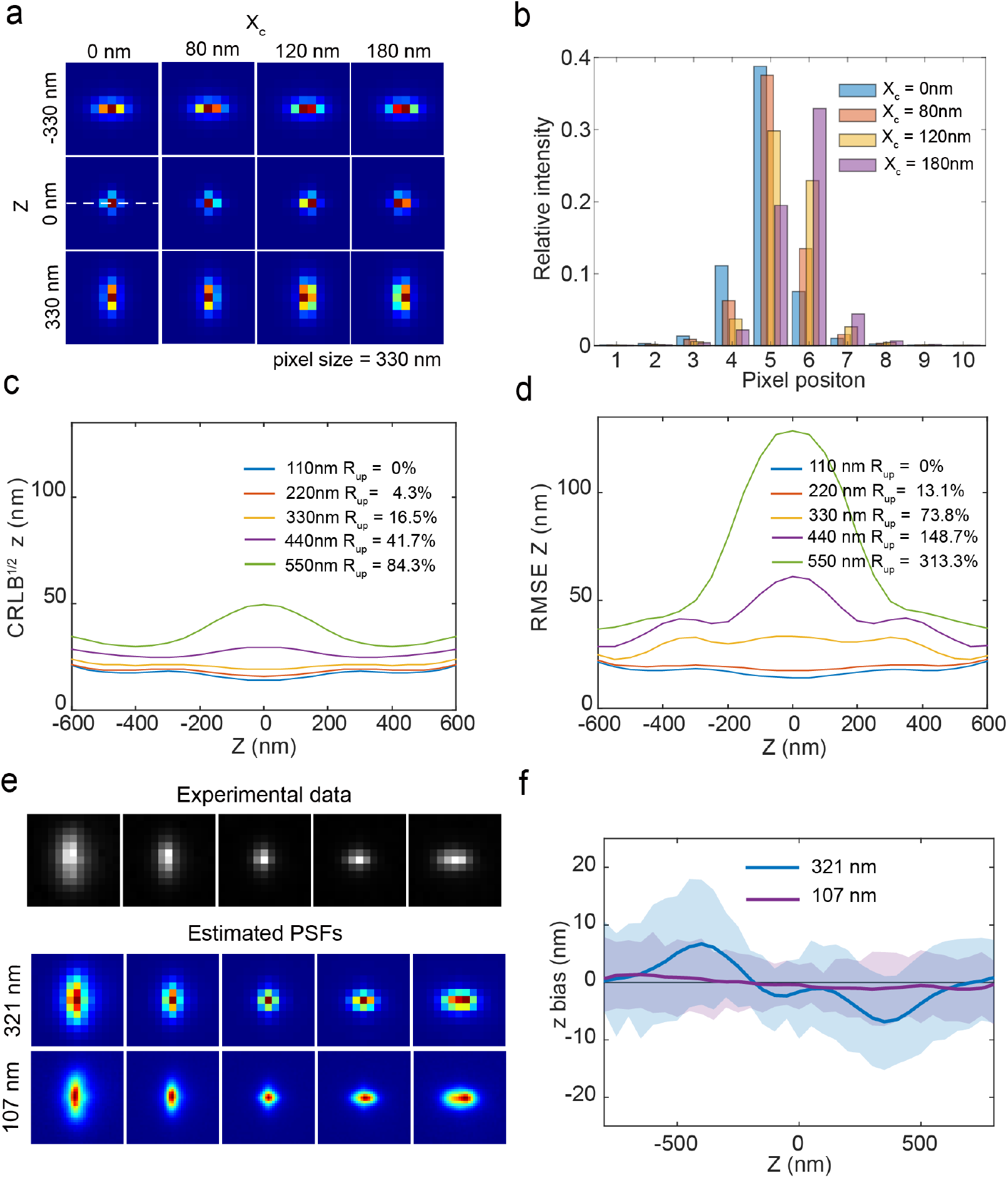
Comparison of CRLB and localization accuracy for PSFs with different pixel sizes. (a) PSFs with a pixel size of 330 nm at different x positions showing the shape of PSF is spatially variant. xc is the x position of the PSF with 0 nm locating at the center of the middle pixel. (b) Relative intensity distribution of the white dashed line in (a) for PSFs at different x positions. (c) Comparison of theoretical axial CRLB for PSFs with different pixel sizes. Each CRLB is averaged from 121 lateral positions uniformly sampled in one pixel. (d) Comparison of axial RMSE for the localization of pixelated simulated data with different pixel sizes using PSFs without up sampling. *R*_*up*_ *rep*resents the percentage degradation in CRLB or RMSE averaged over the range from −600 nm to 600 nm, relative to the values obtained with 110 nm pixel size data. (e) Comparison of the 107 nm and 321 nm pixel size PSFs estimated from the 321 nm pixel size experimental data. (d) Comparison of the impact of the estimated 107 nm and 321 nm pixel size PSFs on localization accuracy of experimental datasets. Solid lines and shaded areas in localization plots indicate mean and standard deviation of the x, y and z bias over >50 beads, respectively.

To validate the accuracy of the up sampled PSF, we conducted a series of evaluations using both simulated and experimental data. First, a fine sampled vector PSF model (SI Note 3) of 22 nm pixel size was employed to accurately represent the continuous PSF. These simulated datasets were then binned using a 15 *×* 15 *pix*el aggregation to produce pixelated PSF data of 330 nm pixel size, mimicking the effects of camera pixelation. Next, from these 330 nm pixel size simulated data, we derived an up sampled PSF with a pixel size of 110 nm. Finally, we utilized the estimated up sampled PSF to re-localize the simulated dataset. As shown in ED Fig 4, single molecule localization with 110 nm pixel size PSF could achieved unbiased and optimal localization accuracy in all 3D, while single molecule localization with 330 nm pixel size PSF generated above results in large localization bias.

Next, we validated the accuracy of the up sampled PSF based localization using experimental data. To this end, we set up a microscope which has a pixel size of 321 nm for the images acquired by the camera to balance between resolution and FOV/ data volume (Methods). We collected many z-stack beads data of 321 nm pixel size randomly distributed at different positions. We then estimated a PSF model with 107 nm pixel size using these beads data as described before. The estimated up sampled PSF model was then applied to localize the z-stack beads data at various positions. We then plot the retrieved x, y and z position bias in each frame as a function of z positions. Any positional deviation in any direction would indicate a mismatch in the model. Similar to the simulated data, the localization results showed that the original 321 nm pixel size PSF (with bin factor 1, SI Note1) based localization led to significant positional bias, while the up sampled 107 nm pixel size PSF (with bin factor 3, SI Note1) achieved unbiased and precise localization (Figure 2e-f, ED Fig 6).

To further demonstrate the performance of the up sampled PSF based localization on biological samples, we first performed 2×2 binning to conventional 127 nm pixel size SMLM data acquired from our reference standard nucleoporin Nup96^19^. Therefore, the data volume was decreased by 75% and the pixel size was down sampled to 254 nm (Figure 3a). Before down sampling, we could nicely resolve the double ring structure of nuclear pore complex (NPC) which separated by only ∼ 50 nm using spline PSF fitter. After down sampling, the conventional spline PSF fitter could not resolve the ring structure with the 254 nm pixel PSF averaged from the down sampled beads data (254 nm pixel size) anymore (Figure 3b). We then derived the 127 nm pixel size PSF from the down sampled beads data. Combined with the up sampled spline fitting algorithm, the 127 nm pixel size PSF based localization could nicely resolve double ring structure from the down sampled single molecule data, indicating that up sample PSF localization could notably improve the accuracy of sparse sampled single molecule data.

**Figure 3.**
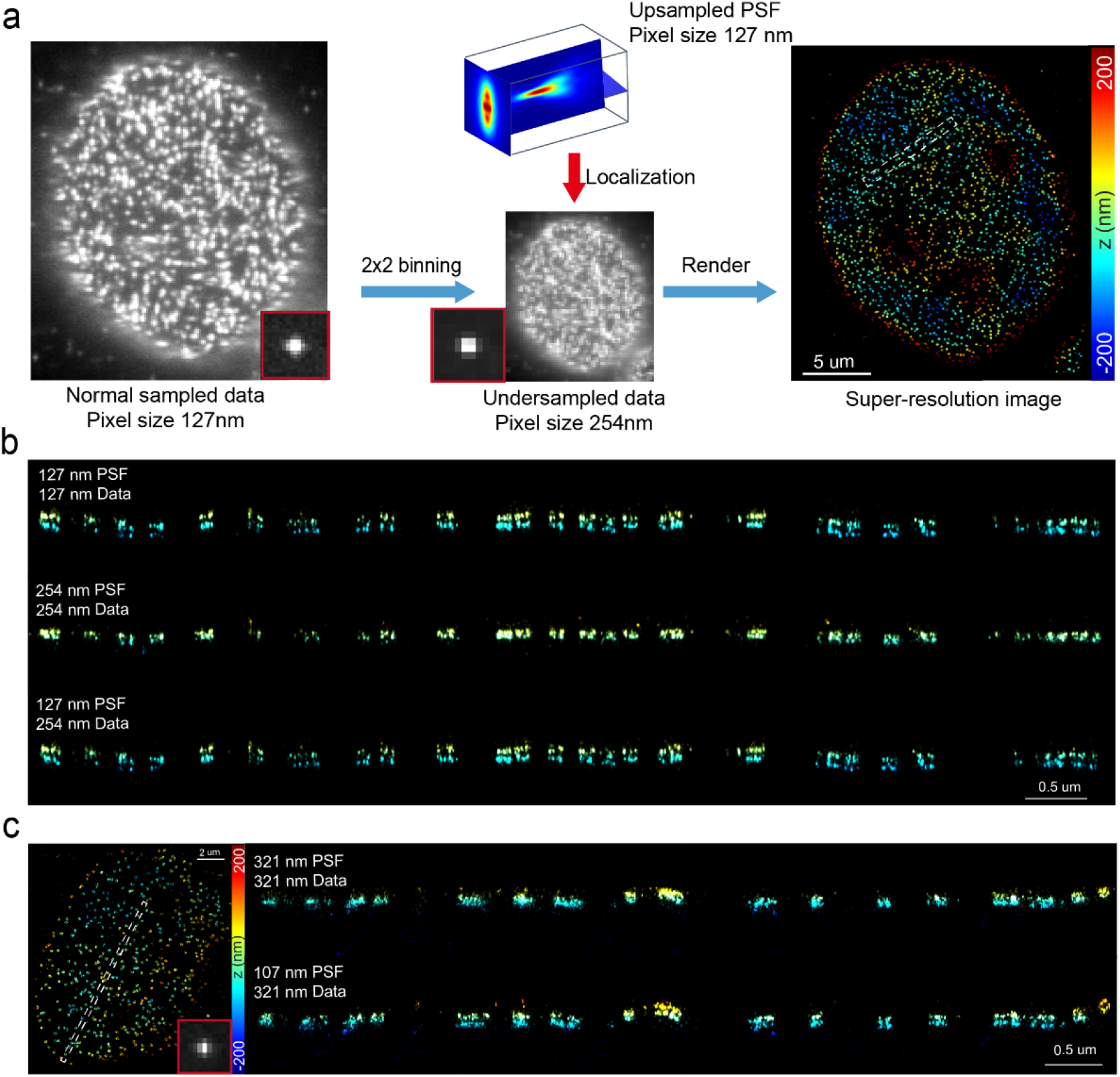
Nup96-AF647 in U2OS cells reconstructed using the up sampled PSF model. (a) Single-molecule data of Nup96-AF647 with a pixel size of 127 nm were 2×2 binned to produce under sampled data with a pixel size of 254 nm. Localization of the under sampled data using the up sampled PSF yielded a super-resolved image. The red boxes indicate the single-molecule image with different pixel sizes. (b) XZ view of the selected region (white dashed lines) in (a) reconstructed from single-molecule data using PSFs of different pixel sizes. (c) Super resolution image of 321 nm pixel size single-molecule data of Nup96-AF647 analyzed with 321 nm and 107 nm pixel size PSFs respectively. The XZ views of the selected region (white dashed lines in the left image of c) were compared.

After showing the effectiveness of the up sampled PSF based single molecule localization, we further increased the pixel size of the acquired image using a microscope (ED Fig 7) with a pixel size of 321 nm. The up sampled PSF model (107 nm pixel size) was estimated from the beads data as described before. As shown in Figure 3c, the double ring structure can’t be resolved if conventional spline fitter was used, while up sampled spline fitter combined with up sampled PSF model could nicely resolve the two rings structure. These findings strongly support the superiority of the up sampled PSF in large-pixel imaging, particularly for applications requiring high-content images with limited pixels and data size compared to the size of the sample (e.g., pathological samples, neuron).

In summary, we proposed an up sampled PSF inverse modeling method to overcome the limitations of large-pixel imaging by recovering high-frequency details and mitigating the trade-offs between pixel size and field of view (FOV). We demonstrated that up sample PSF model could correct the localization bias introduced by pixel integration noise, thus increasing the pixel size by ∼ 2-3 times without losing much resolution. This method not only enhances data accessibility and management but also inspires a rethinking of sampling rates across various biological imaging applications. Accurate up sampled PSF modeling could improve techniques like deconvolution and image denoising, and its potential extends to other fields such as astronomy, where large FOV imaging is critical. By providing a means to achieve both high precision and reduced data burden, this approach could catalyze broader innovations in imaging technologies.

## Methods

### Microscopes

SMLM imaging was performed at room temperature using a custom-built microscope setup (SI Fig. 7). The sample was illuminated via a laser box coupled to a single-mode fiber. The excitation laser was reflected by a dichroic mirror (Di01-R405/488/561/635, Semrock) and focused on the back focal plane of a high numerical aperture (NA) objective (NA 1.5, UPLAPO 100XOHR, Olympus) for super-resolution imaging. Fluorescence emission was collected by the objective and filtered through a quad-band emission filter (NF03-405/488/561/635E, Semrock). Following the tube lens (SWTLU-C, f = 180 mm, Olympus), a band-pass filter (ET685/70m, Chroma) was inserted to remove residual laser light. Subsequently, the fluorescence passed through a 4f system composed of lenses L3 (AC254-150-A-ML, f=150mm, Thorlabs) and L4(AC254-030-A-ML, f=30mm, Thorlabs). Finally, images were acquired using an sCMOS camera (ORCA-Flash4.0 V3, Hamamatsu) with a pixel size of 6.5 *µm* × *6*.*5 µm*. Additionally, a plano-convex cylindrical lens (LJ1516L1-C, Thorlabs) was positioned in front of the camera to introduce astigmatism for 3D imaging.

### Sample preparation

#### Cell Culture

U2OS cells (Nup96-SNAP, catalog no. 300444, Cell Line Services) were cultured in DMEM supplemented with 10% (v/v) fetal bovine serum (FBS), 1x penicillin-streptomycin (PS), and 1x MEM non-essential amino acids (NEAAs; catalog no. 11140-050, Gibco). The cells were maintained at 37°C in a humidified incubator with 5% CO2 and were passaged every 2–3 days. Prior to seeding, 25 mm high-precision round glass coverslips (no. 1.5H, catalog no. CG15XH, Thorlabs) were thoroughly cleaned by sequential sonication in 1 M potassium hydroxide (KOH), Milli-Q water, and ethanol, followed by 30 minutes of UV sterilization. For super-resolution imaging, the U2OS cells were seeded onto the cleaned coverslips and cultured for 2 days until reaching 80-90% confluency. Routine mycoplasma tests confirmed the absence of contamination throughout the study.

#### Fluorescence bead sample

100 nm TetraSpeck beads (Thermo Fisher Scientific, cat. no. T7279) were used. A clean 25 mm coverslip was incubated with 40 µl of 100 mM MgCl2 and 360 µl of 1:1000 diluted bead solution for 5 minutes. The coverslip was then thoroughly rinsed with Milli-Q water three times. After washing, the coverslip was placed in a custom sample holder, and 1 ml of Milli-Q water was added to maintain hydration.

#### Nup96-SNAP-AF647 in U2OS cells

To label Nup96, U2OS-Nup96-SNAP cells were prepared following established protocols. Cells were initially prefixed in 2.4% paraformaldehyde (PFA) for 30 seconds, permeabilized with 0.4% Triton X-100 for 3 minutes, and then fixed in 2.4% PFA for an additional 30 minutes. Following fixation, the cells were quenched with 0.1 M NH4Cl for 5 minutes and washed twice with PBS. To minimize non-specific binding, cells were blocked for 30 minutes using Image-iT FX Signal Enhancer (Invitrogen, catalog no. I36933). For labeling, cells were incubated for 2 hours in a dye solution containing 1 µM SNAP-tag ligand BG-AF647 (New England Biolabs, catalog no. S9136S), 1 mM dithiothreitol (neoFroxx, catalog no. 1111GR005), and 0.5% bovine serum albumin (BSA) in PBS. After incubation, cells were washed three times with PBS for 5 minutes each to remove unbound dye. Finally, cells were post-fixed with 4% PFA for 10 minutes, washed three times with PBS, and stored at 4 C until imaging.

### Data acquisition

#### Collection of bead data on single objective SMLM systems

The fluorescence bead sample was prepared as described above. 30-60 bead stacks were collected where each bead stack was acquired by moving the sample stage from −1 µm to 1 µm with a step size of 50 nm. One frame per z position was collected at an exposure time of 100 ms.

#### Imaging of Nup96-AF647 on single objective SMLM systems

Nup96-SNAP-AF647 labeled U2OS cells were imaged with the Hamamatsu sCMOS camera over an FOV covering 56×56 pixels. The camera was operated under the rolling shutter readout mode with an exposure time of 20 ms. 150,000 frames were acquired. The position of the cylindrical lens before camera was adjusted so that ∼80 nm astigmatism aberration was introduced to the system.

## Data availability

The data that support the findings of this study are publicly available at Zenodo. Data for Fig. 2 are available at https://zenodo.org/records/14000637. Other datasets are available from the corresponding author upon reasonable request.

## Code availability

Up sampled PSF modelling software is available at https://github.com/Li-Lab-SUSTech/Up-sampled-PSF-modeling, it includes source codes, example Jupyter notebooks and tutorials for localization analysis using the up sampled PSF models.

## Extended Figures

**ED Fig 1.**
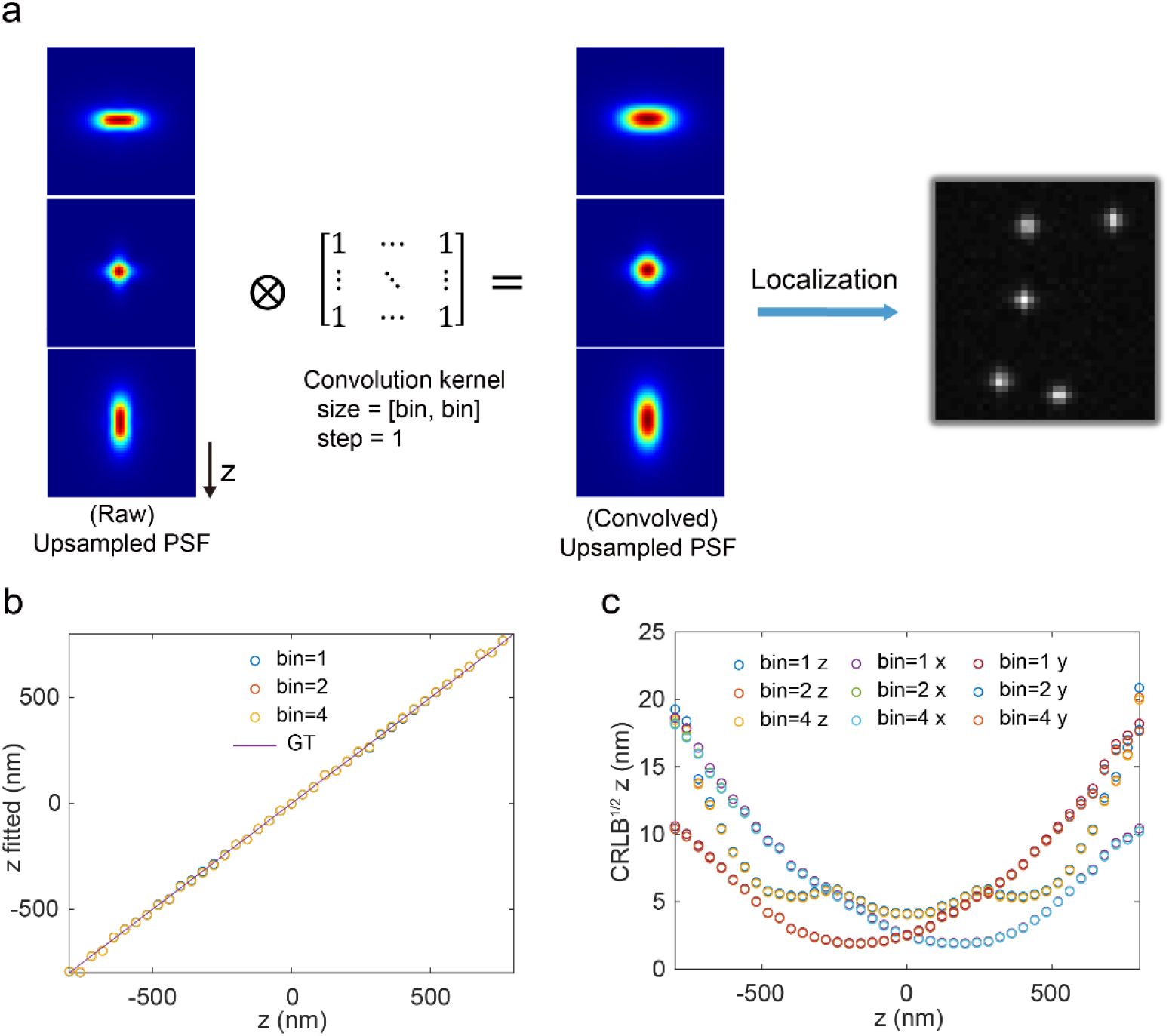
Validation of Spline fitting for different sampling rates. (a) To match the pixel size of the up sampled PSF with the simulated or experimental data, a convolution operation is performed on the estimated raw up sampled PSF (Raw) to generate the pixel integrated up sampled PSF (Convolved), using a convolution kernel size of bin × bin with all values set to 1 (SI Note 2 Eqs. 2.1). The convolved PSF is used for localizing experimental data. The *bin* factor represents the ratio between the pixel size of the simulated or experimental data and the pixel size of the up sampled PSF, with its value constrained to integer values. (b) The spline fitter returns the z-position results by fitting the same simulated data using up sampled PSFs at different sampling rates. (c) Localization results of 110, 55 and 27.5 nm pixel size PSF for 110 nm pixel size single molecule data. The simulated data consists 51 axial positions evenly distributed along −600 nm to 600 nm with 1,000 single molecules at each axial position. Each molecule contributes a total of 2,000 photons, with a background level of 2 × 10^−3^ *photons/nm*^2^.

**ED Fig 2.**
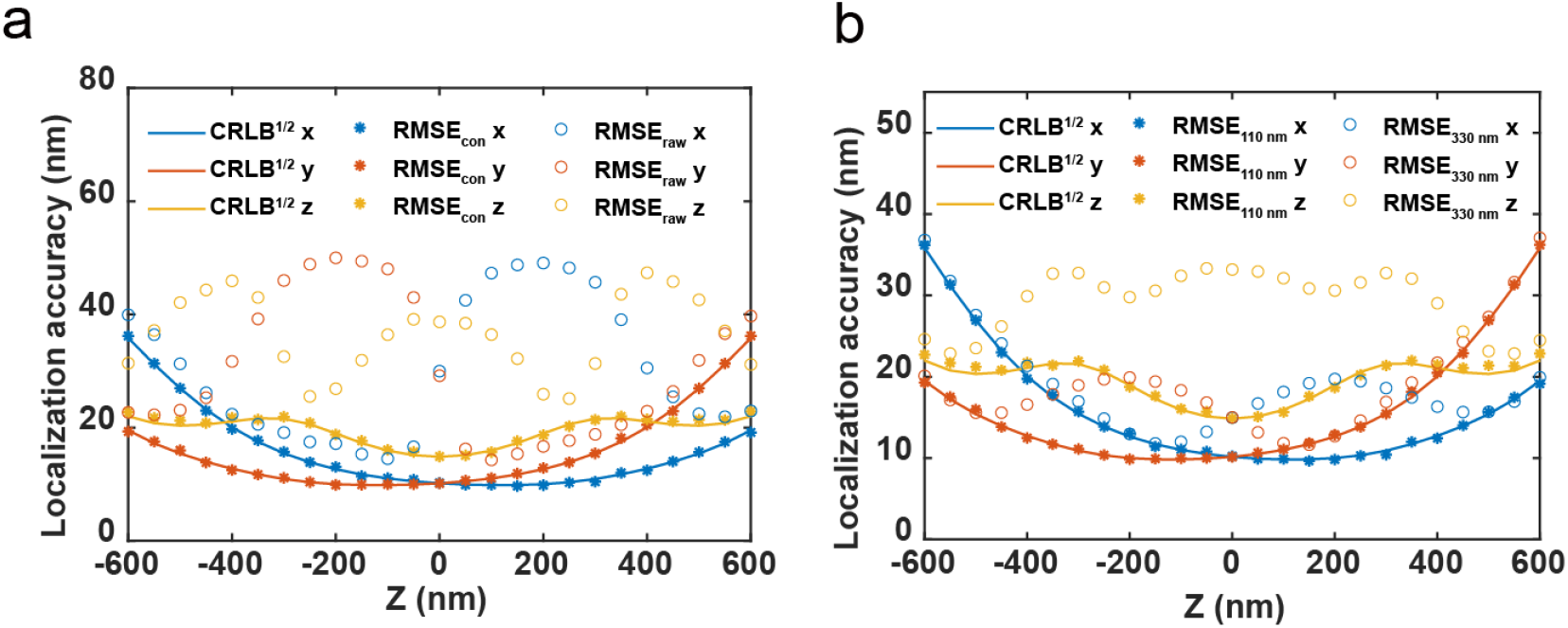
CRLB and RMSE improved by using up sampled PSF model and spline PSF fitter. (a) Comparison of RMSE with and without considering pixel integration for data with pixel size of 330 nm. RMSEcon represents the localization results of the up sampled PSF (110nm) after convolution, while RMSEraw represents the localization results of the up sampled PSF (110nm) without convolution. (b) Comparison of the RMSE for localizing the 330 nm pixel size data using 110 nm PSF and 330 nm PSF. The simulation parameters are the same as those in ED Fig 1.

**ED Fig 3.**
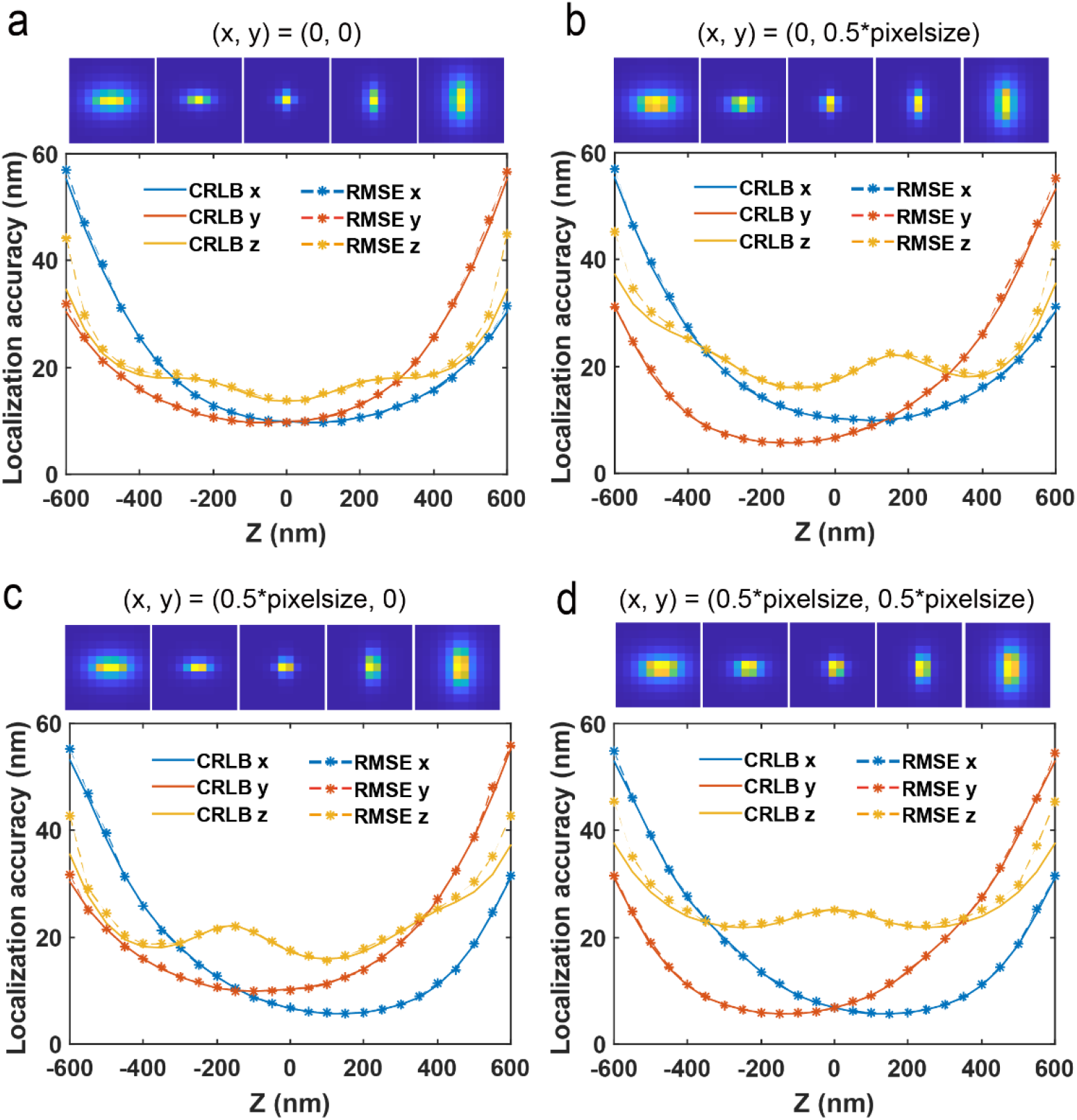
Comparison of localization accuracy at different lateral positions. Localization accuracy of 110 nm pixel size PSFs on 330 nm single molecule data at different lateral positions, (0 nm, 0 nm), (0 nm, 115 nm), (115 nm, 0 nm), and (115 nm, 115 nm) for a, b, c, and d respectively. Here, the center of the central pixel is set as coordinate (0 nm, 0 nm). We used a vector PSF model (SI Note 3) to generate 1,000 simulated single molecules at different x and y positions at each axial position. 25 axial positions uniformly distributed along −600 nm to 600 nm were evaluated. Each molecule contributes a total of 2,000 photons, with a background level of 2 × 10^−3^ *photons/nm*^2^. Localization accuracy at each z position is computed from the root mean square error (RMSE) of the localization results from 1,000 single molecules.

**ED Fig 4.**
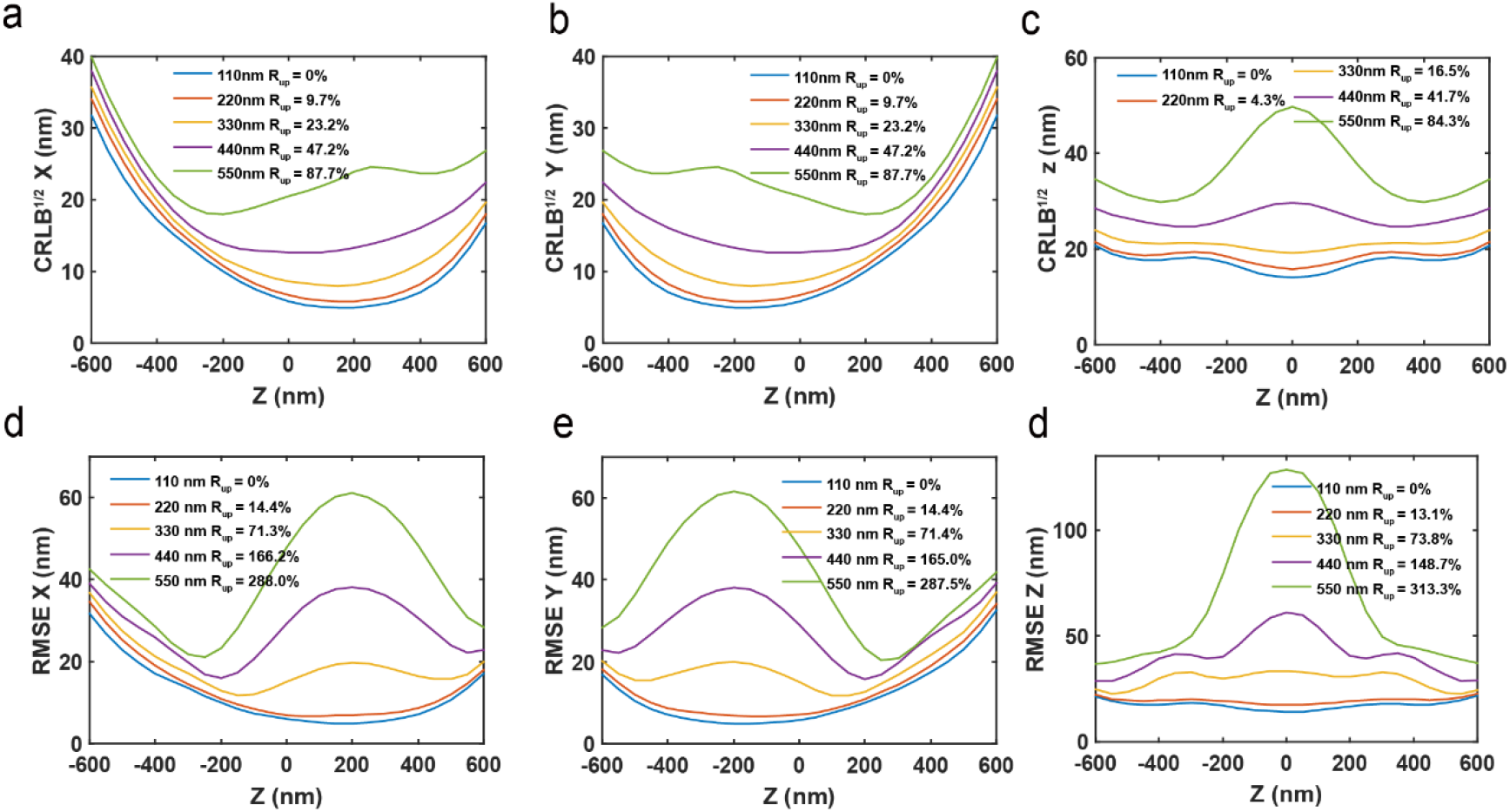
Comparison of the CRLB and RMSE for PSFs with different pixel sizes. (a-c) Comparison of the theoretical CRLB for x, y, and z for PSFs with different pixel sizes. (d-f) Comparison of the RMSE in the x, y, and z directions localized with PSFs of different pixel sizes. *R*_*up*_ represents the percentage degradation in CRLB or RMSE averaged over the range from −600 nm to 600 nm, relative to the values obtained with a pixel size of 110 nm. The simulation parameters are the same as those in ED Fig 3.

**ED Fig 5.**
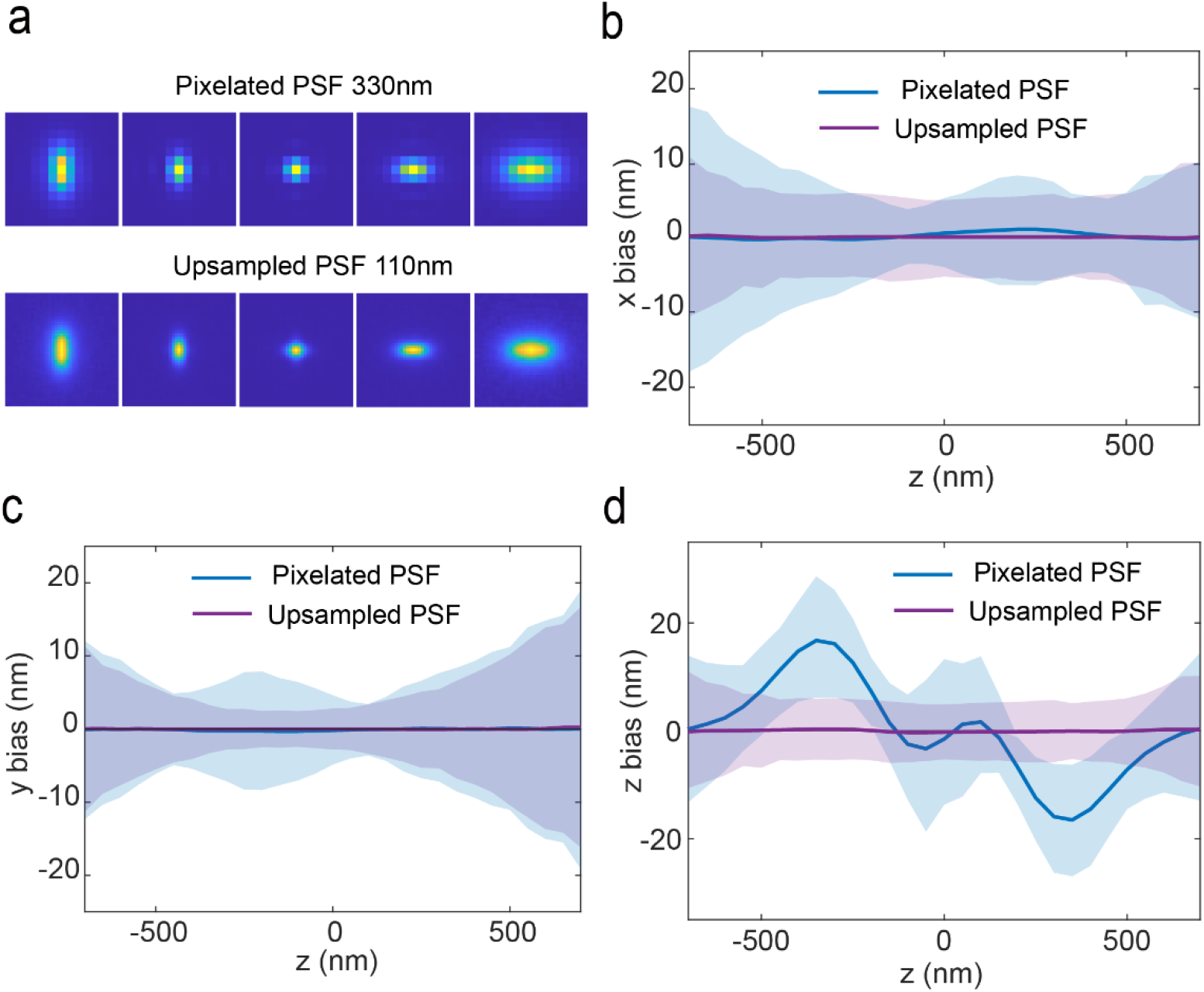
Validation of the up sampled PSF modelling using simulated data. (a) The 330 nm pixel size PSF (top) and 110 nm pixel size PSF (bottom) estimated from 300 simulated data sets with a pixel size of 330 nm. (b-d) Comparison of the impact of the estimated 330 nm and 110 nm pixel size PSF on the localization accuracy of the 330 nm pixel size simulated data along x, y and z directions, respectively. Each molecule contributes a total of 10,000 photons, with a background level of 2 × 10^−3^ *photons/nm*^2^. The x/y/z bias as a function of z are defined as in SI Note 5 Eqs. 5.2 and 5.1. Solid lines and shaded areas in localization plots indicate mean and standard deviation of bias, respectively.

**ED Fig 6.**
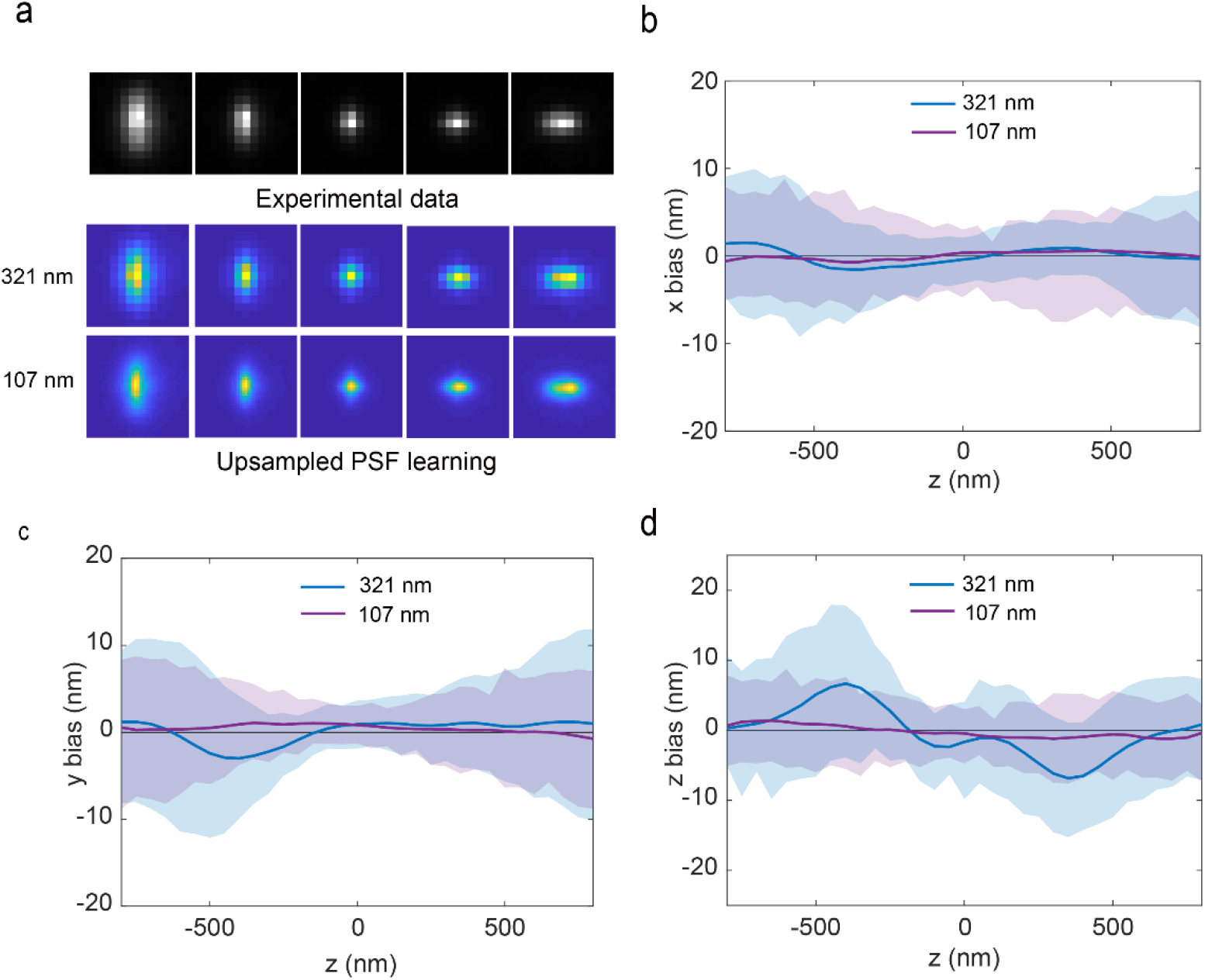
Validation of the up sampled PSF modelling using experimental data. (a) 321 nm pixel size experimental bead data (top) and the estimated 321 nm and 107 nm pixel size PSF (bottom). (b-d) Comparison of the impact of the estimated 321 nm and 107 nm pixel size PSF on the localization accuracy of the 321 nm pixel size experimental data along x, y and z directions, respectively. The x/y/z bias vs z are defined as in SI Note 5 Eqs. 5.2 and 5.1. Solid lines and shaded areas in localization plots indicate mean and standard deviation of bias over >50 beads, respectively.

**ED Fig 7.**
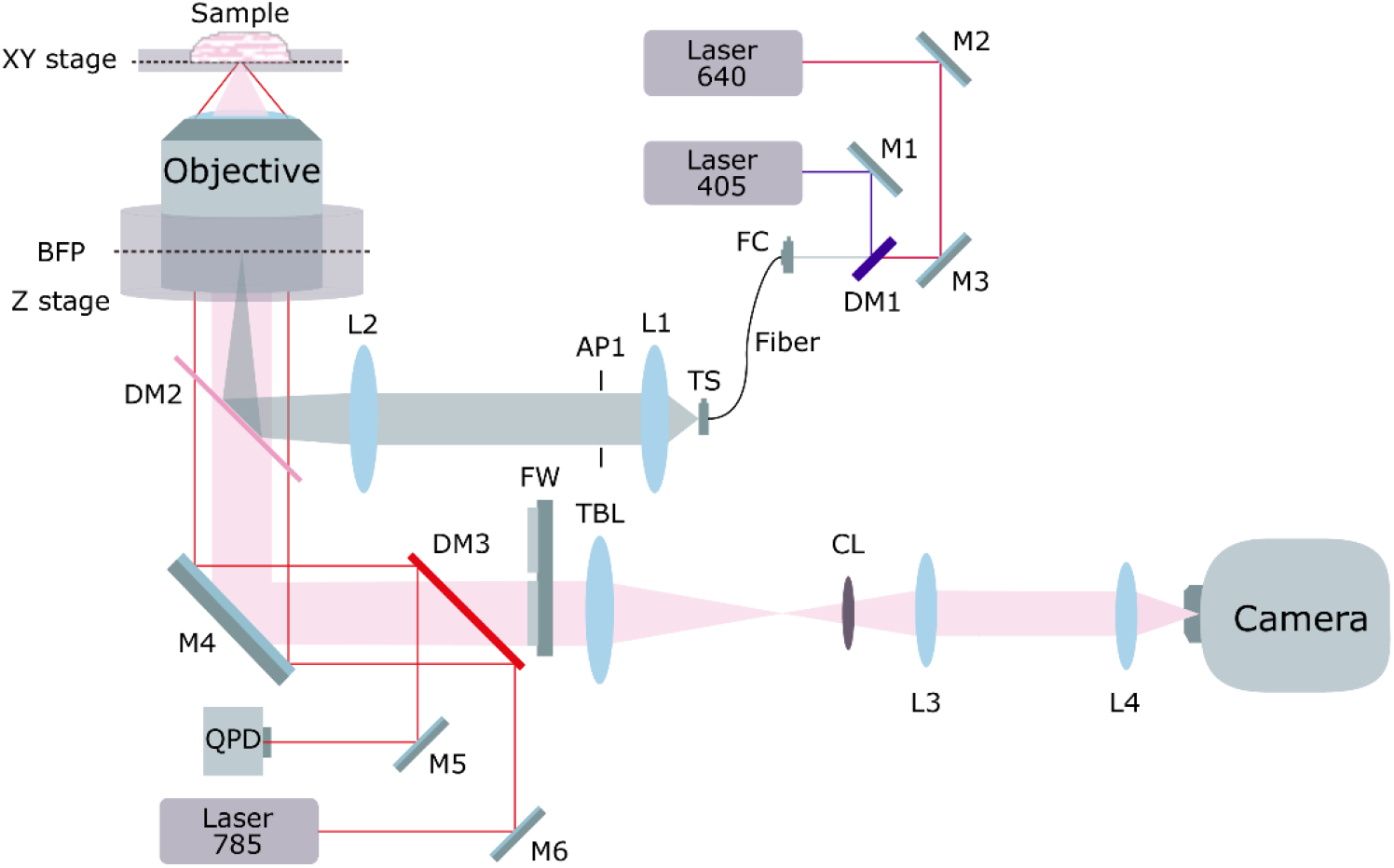
Detailed layout of the optical setup. M, mirror; DM, dichroic mirror; L, lens; TS, translation stage; FC, fiber coupler; Fiber, single-mode fiber; BFP, back focal plane; FW, filter-wheel; TBL, tube lens; AP, aperture; QPD, quadrant photodiode. The excitation lasers are first reflected by dichroic mirror DM1 and coupled into a single-mode fiber through the fiber coupler FC. Before being reflected by the main dichroic mirror DM2 to enter the objective for sample illumination, the beam is collimated and reshaped by a pair of lenses (L1 and L2) and a slit at AP1. In the imaging path, the fluorescence collected by the objective passes through the dichroic mirror DM2 and is filtered by the filter wheel FW. It is then focused using a tube lens TBL. Subsequently, the fluorescence passes through a 4f system composed of lenses L3 (focal length =150nm) and L4 (focal length =30nm), before being detected by the camera. Additionally, a beam excited by a 785 nm laser, reflected off the coverslip, is detected by the quadrant photodiode (QPD), providing feedback control to the z-stage for focus locking.

## References

1. Schermelleh, L. et al. Super-resolution microscopy demystified. Nat. Cell Biol. 21, 72–84 (2019).

2. Lelek, M. et al. Single-molecule localization microscopy. Nat. Rev. Methods Prim. 1, (2021).

3. Holden, S. J. et al. High throughput 3D super-resolution microscopy reveals Caulobacter crescentus in vivo Z-ring organization. Proc. Natl. Acad. Sci. U. S. A. 111, 4566–4571 (2014).

4. Mahecic, D. et al. Homogeneous multifocal excitation for high-throughput super-resolution imaging. Nat. Methods 17, 726–733 (2020).

5. Beghin, A. et al. Localization-based super-resolution imaging meets high-content screening. Nat. Methods 14, 1184–1190 (2017).

6. Fu, S. et al. Field-dependent deep learning enables high-throughput whole-cell 3D super-resolution imaging. Nat. Methods 20, 459–468 (2023).

7. Barentine, A. E. S. et al. An integrated platform for high-throughput nanoscopy. Nat. Biotechnol. 41, 1549–1556 (2023).

8. Thompson, R. E., Larson, D. R. & Webb, W. W. Precise nanometer localization analysis for individual fluorescent probes. Biophys. J. 82, 2775–2783 (2002).

9. Mortensen, K. I., Churchman, L. S., Spudich, J. A. & Flyvbjerg, H. Optimized localization analysis for single-molecule tracking and super-resolution microscopy. Nat. Methods 7, 377–381 (2010).

10. Chang, H., Fu, S. & Li, Y. Optimal sampling rate for 3D single molecule localization. Opt. Express 31, 39703 (2023).

11. Babcock, H. P. & Zhuang, X. Analyzing Single Molecule Localization Microscopy Data Using Cubic Splines. Sci. Rep. 7, 1–8 (2017).

12. Li, Y. et al. Real-time 3D single-molecule localization using experimental point spread functions. Nat. Methods 15, 367–369 (2018).

13. Liu, S. et al. Universal inverse modeling of point spread functions for SMLM localization and microscope characterization. Nat. Methods 21, 1082–1093 (2024).

14. Laurence, T. A. & Chromy, B. A. Efficient maximum likelihood estimator fitting of histograms. Nat. Methods 7, 338–339 (2010).

15. Liu, T. & Li, D. Convergence of the BFGS-SQP method for degenerate problems. Numer. Funct. Anal. Optim. 28, 927–944 (2007).

16. Stallinga, S. & Rieger, B. Accuracy of the Gaussian Point Spread Function model in 2D localization microscopy. Opt. Express 18, 24461 (2010).

17. Fu, S. et al. Deformable mirror based optimal PSF engineering for 3D super-resolution imaging. Opt. Lett. 47, 3031 (2022).

18. Huang, B., Wang, W., Bates, M. & Zhuang, X. Three-Dimensional Super-Resolution Reconstruction Microscopy. Science, 319, 810–813 (2008).

19. Thevathasan, J. V. et al. Nuclear pores as versatile reference standards for quantitative superresolution microscopy. Nat. Methods 16, 1045–1053 (2019).

