## Supplementary information for "Up sampled PSF enables high accuracy 3D super-resolution imaging with sparse sampling rate"

### Supplementary Notes

#### 1. Up sampled PSF modelling in the spatial domain

##### 1.1 Calculation of initial values

The initial position for each bead is,

$$x_{init} = y_{init} = z_{init} = 0. \tag{1.1}$$

The background value for each bead is estimated as

$$Bg_i = \min\Big(D_i(x,y,z) \otimes G\big(x,y,z,\sigma_x = \sigma_y = \sigma_z = 2\big)\Big)/bin^2, \tag{1.2}$$

Where $D_i(x,y,z)$ represents the cropped 3D image stack of bead $i$, while $G$ denotes a 3D Gaussian kernel with standard deviations $\sigma_x$, $\sigma_y$, and $\sigma_z$ in the respective $x$, $y$, and $z$ directions. $bin$ factor refers to the ratio between the pixel size of the data and the pixel size of the up sampled PSF. The symbol $\otimes$ denotes the convolution operation. The minimum value is taken over all voxels within the cropped and filtered region. The initial background for each bead is set to the median value of $bg_i$ across all bead data,

$$Bg_{init} = \underset{i}{\text{median}}(b_i). \tag{1.3}$$

The initial photon value for each bead is estimated as

$$Np_{init}^{i} = \underset{z}{\text{avg}}\left(\sum_{x,y} D_i(x,y,z) - b_i\right). \tag{1.4}$$

The initial PSF model is a 3D array with each element set to $1/N_s^2/bin^2$ with $N_s$ the ROI size of bead data. For example, if the ROI size is $N_s \times N_s = 21 \times 21$ pixels and the sampling factor $bin$ is 4, the initial value is 0.00014. This assumes that the sum of each axial slice of the PSF model is one.

##### 1.2 Calculation of the forward model using the up sampled PSF

The up sampled PSF model at a given bead location $(x_i,\ y_i,\ z_i)$ can be calculated as follows[1],

$$U_i^{up}(x - x_i, y - y_i, z - z_i) = \mathcal{F}_{3D}^{-1}\big(\mathcal{F}_{3D}\big(Np_i \cdot PSF_{up}(x,y,z) + Bg_i\big) \cdot g(q) \cdot e^{i\varphi_{\text{shift}}}\big), \tag{1.5}$$

Where the shifting phase is calculated from,

$$\varphi_{\text{shift}} = 2\pi\big(q_x x_i + q_y y_i + q_z z_i\big), \tag{1.6}$$

where $q_x, q_y, q_z$ are the Cartesian coordinates of the frequency space as given by,

$$q_x = \frac{x}{L_x}, \qquad q_y = \frac{y}{L_y}, \qquad q_z = \frac{z}{L_z}, \tag{1.7}$$

where $L_x, L_y$ and $L_z$ are length of the up sampled PSF model in $x, y$ and $z$ directions. Considering the bead size, we model the bead as a sphere and $g(q)$ is the analytical Fourier transform of a solid sphere of radius $r_0$,

$$g(q) = \frac{J_{\frac{3}{2}}(2\pi q r_0)}{(q r_0)^{\frac{3}{2}}} r_0^3, \tag{1.8}$$

where $J$ is the Bessel function of the first kind and $q$ is the is the spherical coordinate in the frequency space and is calculated from,

$$q = \sqrt{q_x^2 + q_y^2 + q_z^2}, \tag{1.9}$$

To ensure that the variation of $U_i^{up}$ is not affected, $g(q)$ is normalized to its maximum value. To match $U_i^{up}$ with the pixel size of the data, we further apply a binning operation to $U_i^{up}$,

$$U_i(k,l) = \sum_{i=1}^{bin} \sum_{j=1}^{bin} U_i^{up}(bin \cdot k + i, bin \cdot l + j), \tag{1.10}$$

Where $U_i$ is the binned (merged) forward model, which can be directly compared with the measured bead data $M_i$. $k, l$ denote the horizontal and vertical pixel indices of the forward model.

##### 1.3 Variable scaling

We utilized the L-BFGS-B algorithm from the SciPy optimization package for our optimization process. To account for the varying scales of gradients across different variable types, we applied a tailored scaling factor to each type. This adjustment ensured uniform update rates for all variables throughout each iteration. The original variables in the forward model were substituted with transformed variables, ensuring consistent and balanced updates during the optimization procedure. For example,

$$Np_i = \frac{Np_i}{w_{Np_i}} w_{Np_i} = Np_{w,i} w_{Np},$$

$$Bg_i = \frac{bg_i}{w_{bg_i}} w_{bg_i} = Bg_{w,i} w_{bg}, \tag{1.11}$$

$$PSF_{up} = \frac{PSF_{up}}{w_{PSF_{up}} bin^2} w_{PSF_{up}} = PSF_W w_{PSF},$$

where $PSF_w, Np_{w,i}, bg_{w,i}$ are scaled variables, with respect to which the gradient will be calculated, and $w_{PSF}, w_{Np}, w_{bg}$ are scaling factors for the up sampled PSF model, photons and background. The scaling factors can be determined based on the gradient values of their respective parameters, ensuring that all gradient values are within the same order of magnitude. Proper variable scaling is crucial in PSF learning, as it ensures that the optimization process converges to the global minimum and facilitates the efficient optimization of all variables.

##### 1.4 Calculation of loss function

The loss function for up sampled PSF estimation is,

$$loss = LL + a_{\mathrm{drift}} f_{\mathrm{drift}} + \beta(a_{\mathrm{PSF,min}} f_{\mathrm{PSF,min}} + a_{\mathrm{Bg,min}} f_{\mathrm{Bg,min}} + a_{\mathrm{Np,min}} f_{\mathrm{Np,min}} + a_{\mathrm{norm}} f_{\mathrm{norm}}), \tag{1.12}$$

Where $LL$ is based the log-likelihood function, assuming that the converted pixel values follow a Poisson distribution,

$$LL = \underset{i}{\text{avg}}[U_i - D_i - D_i \log(U_i) + D_i \log(D_i)], \tag{1.13}$$

Note that $D_i$ and $D_i \log(D_i)$ are normalization factors for the likelihood probabilities, aimed at improving the stability of the fitting process. When sample drift is considered, $f_{\text{drift}}$ is equal to the $L^1$ norm of all drift rates over the bead data, this term is to constrain the estimated drift rates to be close to zero, to avoid adding an arbitrary constant drift rate.

The next three terms serve to constrain the values of up sampled PSF model, photons and background to be positive,

$$\begin{aligned} f_{\text{PSF,min}} &= \sum_{x,y,z} \min\left(PSF_{up}, 0\right)^2, \\ f_{\text{Np,min}} &= \sum_{i} \min(Np_i, 0)^2, \\ f_{\text{Bg,min}} &= \sum_{i} \min(Bg_i, 0)^2 \end{aligned} \tag{1.14}$$

Here min denotes element-wise minimum comparing each value in an array with zero. As default we set $a_{\text{PSF,min}} = 100, a_{\text{Np,min}} = 1, a_{\text{Bg,min}} = 1$.

#### 2. Localization Methods Using Up sampled PSF

Before performing cubic spline interpolation on the estimated up sampled PSF, a convolution operation must first be applied. This is because the experimental data are obtained through integration over large pixel sizes, while the estimated up sampled PSF is based on integration over small pixel sizes. The difference between the two is determined by the $bin$ factor, which represents the ratio between the large and small pixel sizes. To align the two, a convolution is performed on the upsampled PSF using a kernel of size corresponding to the $bin$ factor, with all elements set to 1, and a stride of 1. The detailed convolution procedure is as follows,

$$PSF_{con}(k,l) = PSF_{raw} \otimes ones(bin, bin) = \sum_{i=1}^{bin} \sum_{j=1}^{bin} PSF_{raw}(s*k+i, s*l+j) \tag{2.1}$$

where $PSF_{con}$ represents the convolved up sampled PSF, $\otimes$ denotes the convolution operation, and $ones(bin)$ is a matrix of size equal to the $bin$ factor, with all elements set to 1. The $s$ denotes the convolution stride, fixed at 1 in this instance. The $bin$ factor refers to the ratio between the pixel size of the data and the pixel size of the upsampled PSF.

For the localization step using the $PSF_{con}$, the data was analyzed with the cubic spline fitting method. The cubic spline interpolation of a given PSF model is[4,5],

$$f_{i,j,k}(x,y,z) = \sum_{m=0}^{3} \sum_{n=0}^{3} \sum_{p=0}^{3} a_{i,j,k,m,n,p} \left( \frac{x - x_i}{\frac{\Delta x}{bin}} \right)^m \left( \frac{y - y_j}{\frac{\Delta y}{bin}} \right)^n \left( \frac{z - z_k}{\Delta z} \right)^p \tag{2.2}$$

where $\Delta x$, $\Delta y$ are the x and y pixel sizes of the camera, $\Delta z$ is the axial step size of the z-stack, $a_{i,j,k,m,n,p}$ are the spline coefficients and $x_i, y_j, z_k$ are the start position of each voxel $(i,j,k)$ in the up sampled PSF. $x, y$ are the positions corresponding to the camera pixels, while $z$ represents the position in the z-stack. After building the spline PSF model, maximum likelihood estimation (MLE) with Poisson statistics was used to localize beads or single molecules with the objective function given by[6],

$$\chi^2_{mle} = 2\left(\sum_{k,j}(U_{k,j} - M_{k,j}) - \sum_{k,j,M_{k,j}>0} M_{k,j} \ln\frac{U_{k,j}}{M_{k,j}}\right) \tag{2.3}$$

where $U_{k,j}$ and $M_{k,j}$ are the expected photon number and measured photon number in the pixel $(k, j)$, respectively. We used a modified Levenberg-Marquardt (L-M) algorithm[7] to minimize $\chi^2_{mle}$ for the parameter estimation.

### 3. Vectorial PSF model calculation

To accurately model the image formation process in a microscope equipped with a high numerical aperture (NA) objective, a vectorial PSF model is employed, accounting for the refractive index mismatch between the medium-cover slip interface and the cover slip-immersion medium interface. Given that fluorescent probes are typically flexibly attached to the target molecules and are capable of free rotation, we assume an isotropic emitter PSF model for our analysis. The vectorial PSF can be expressed as[2]:

$$PSF(x - x_i, y - y_i, z - z_i) \propto \sum_{\substack{m=x,y \\ n=p_x,p_y,p_z}} \left|\mathcal{F}_{czt}\left\{h(k_x, k_y) w_{mn} e^{i2\pi[k_x(x-x_i)+k_y(y-y_i)-k_z(z-z_i)]}\right\}\right|^2 \tag{3.1}$$

$$k_z = \sqrt{k^2 - k_x^2 - k_y^2}$$

where $h(k_x, k_y)$ is pupil function and can be written as,

$$h(k_x, k_y) = T_a A(k_x, k_y) e^{i\Phi(k_x,k_y)}, \tag{3.2}$$

where $A$ and $\Phi$ are the magnitude and phase components of the pupil function, each of them is a 2D array. $T_a$ is the apodization factor of the objective lens and is equal to

$$T_a = \frac{\sqrt{cos\theta_{imm}}}{cos\theta_{med}}, \tag{3.3}$$

where $\theta_{\text{med}}$ and $\theta_{\text{imm}}$ represent the angles of the optical rays in the sample medium and the immersion medium, respectively, and are constrained by the numerical aperture $(NA)$ of the objective lens. The wave vector $k$ has Cartesian components $k_x, k_y, k_z$, with its magnitude$k = \frac{n_{imm}}{\lambda}$, where $n_{imm}$ is the refractive index of the immersion medium, and $\lambda$ is the central wavelength corresponding to the emission filter. The term $e^{-i2\pi k_z(z-z_i)}$ accounts for the defocus phase in the propagation.

We employed the Chirp Z-transform based on Bluestein's algorithm (denoted as $\mathcal{F}_{czt}$) to compute the 2D Fourier transform of the pupil function. The advantage of using Bluestein's algorithm is that the pupil function size becomes independent of the camera's pixel size, allowing a pupil size of 64 × 64 pixels to offer sufficient sampling for accurate representation.

In this context, $w_{mn}$ denotes the mmm-component of the electric field at the pupil plane generated by the $n$-component of the dipole moment of the fluorophore. The calculation of $w_{mn}$ is as follows,

$$\begin{aligned} w_{xn} &= P_n cos\varphi - S_n sin\varphi, \\ w_{yn} &= P_n sin\varphi - S_n cos\varphi, \end{aligned} \tag{3.4}$$

where $\varphi$ is the angular component in the polar coordinate of the frequency space. $P_n$ and $S_n$ are electric field components in p and s polarizations relative to the incident plane at the sample space,

$$P_{px} = T_p cos\theta_1 cos\varphi,$$

$$P_{py} = T_p cos\theta_1 sin\varphi,$$

$$P_{pz} = -T_p sin\theta_1, \tag{3.5}$$

$$S_{pz} = -T_s sin\varphi,$$

$$S_{py} = T_s cos\varphi,$$

$$S_{pz} = 0,$$

where $p_x, p_y, p_z$ are the Cartesian components of the dipole moments, $T_P$ and $T_S$ are the total transmission coefficients of p- and s-polarized light. The fluorescence light starting from the dipole emitter propagates through the sample medium, the coverslip and the immersion medium. In this three-layer system, we ignore the multiple reflections at the two interfaces, then $T_P$ and $T_S$ can be calculated from,

$$T_p = \tau_{P13}\tau_{P23},$$
$$T_s = \tau_{S13}\tau_{S23}, \tag{3.6}$$

where $\tau_{Pij}$ and $\tau_{Sij}$ are the Fresnel transmission coefficients of s- and p-polarized light travel from medium $i$ to medium $j$,

$$\tau_{Pij} = \frac{2n_i \cos\theta_i}{n_i \cos\theta_j + n_j \cos\theta_i},$$
$$\tau_{Sij} = \frac{2n_i \cos\theta_i}{n_i \cos\theta_i + n_j \cos\theta_j}, \tag{3.7}$$

where $n_i$ and $\theta_i$ are the refractive index and the light propagation angle in medium $i$, and the subscript 1,2,3 denotes the sample medium, the coverslip and the immersion medium respectively.

In order to realistically account for the effects of fluorescence dipole emission, pixelation, particle size, dispersion, and chromatic aberration, the PSF model is rescaled using the OTF as,

$$PSF_{otf} = \mathcal{F}_{3D}^{-1}[\mathcal{F}_{3D}(PSF) \cdot e^{-i2(\sigma_x q_x \pi)^2 - i2(\sigma_y q_y \pi)^2}], \tag{3.8}$$

where $\sigma_x$ and $\sigma_y$ are the standard deviation (in pixel unit) of a 2D Gaussian kernel in real space. By including the photon count and background of each bead stack, the final forward model is,

$$U = Np \cdot PSF_{otf} + bg \tag{3.9}$$

Where $Np$ and $bg$ represent the photon count and background, respectively.

#### 4. Analytical CRLB Calculation for Pixelated PSF

The CRLB is calculated from the diagonal elements of the inverse of the Fisher information matrix $I(\theta)$, which measures the amount of information that an observation (PSF) carries about the estimated parameters $\Theta$.

$$CRLB(\theta)_{ij} = diag(I(\theta)_{ij}^{-1}) \tag{4.1}$$

Where $diag$ refers to extracting the diagonal elements of a matrix. The Fisher information matrix is defined as,

$$I(\theta)_{ij} = \sum_j \sum_k \frac{1}{U_k} \frac{\partial U_{k,l}}{\partial \theta_i} \frac{\partial U_{k,l}}{\partial \theta_j}, with\ U_{k,l} = [Np \cdot PSF_{pixelated}(k,l) + bg]_k \tag{4.2}$$

where $\Theta$ is a set of parameters being estimated, $k, l$ denote the horizontal and vertical pixel indices of the pixelated PSF, respectively. The parameters $\theta$ for pixelated PSF is $x, y, z$, photon $Np$, and background $bg$ for each emitter. The $U_{k,l}$ is the forward model as defined in Eq. 1.10.

The pixelated PSF and its first-order derivatives with respect to the parameters are obtained by integrating the up sampled PSF and its corresponding derivatives. The first-order derivatives of the pixelated forward model $U_k$ with respect to the parameters $\Theta$ are given by,

$$PSF_{pixelated}(k,l) = \sum_{i=1}^{bin} \sum_{j=1}^{bin} PSF_{upsampled}(bin \cdot k + i, bin \cdot l + j) \tag{4.3}$$

$$\frac{\partial U_{k,j}}{\partial \theta_{x,y,z}} = Np \sum_{i=1}^{bin} \sum_{j=1}^{bin} \sum_{\substack{m=x,y \\ n=p_x,p_y,p_z}} Re\left\{E_{mn}^* \frac{\partial E_{mn}}{\partial \Theta_{x,y,z}}\right\}$$

$$\frac{\partial U_k}{\partial Np} = PSF_{pixelated}(k,l)$$

$$\frac{\partial U_k}{\partial bg} = 1$$

$$E_{mn} = \mathcal{F}_{\mathrm{czt}}\left[h(k_x, k_y) w_{mn} e^{i2\pi(k_{z\mathrm{med}} z_i - k_z z_s)} e^{i2\pi(k_x x_i + k_y y_i)}\right]$$

$$\frac{\partial E_{mn}}{\partial \Theta_{x,y,z}} = \mathcal{F}_{\mathrm{czt}}\left[i k_{x,y,z\mathrm{med}} h(k_x, k_y) w_{mn} e^{i2\pi(k_{z\mathrm{med}} z_i - k_z z_s)} e^{i2\pi(k_x x_i + k_y y_i)}\right]$$

where $bin$ factor refers to the ratio between the pixel size of the data and the pixel size of the upsampled PSF.

Due to the significant variation in CRLB at different positions within the large-pixel vector PSF (ED Fig. 3), 121 points were uniformly selected within a single pixel. The CRLB was calculated for each point, and the average of these values was taken to obtain the final mean CRLB.

$$CRLB_{mean}^{\frac{1}{2}} = \sum_{i=1}^{num} CRLB^{\frac{1}{2}}(x_i, y_i) / num \tag{4.4}$$

Where $x_i$ and $y_i$ represent different position coordinates within a single pixel. The $num$ represents the number of different positions sampled within a single pixel.

The CRLB calculated in current work is slightly better than that in previous work using numerical approach, especially for lager pixel conditions. We found that blurring of the theoretical PSF with a Gaussian function (Eqs. 2.8) to mimic experimental PSF will reduce the impact of the pixel size on the theoretical CRLB, which was not considered in Huang et al. (SI Fig 1). We also showed that our up sampled spline PSF fitter could achieve a localization accuracy approaching the calculated CRLB (ED Fig. 3), showing that both the up sampled PSF modeling and up sampled spline PSF fitter could achieve the optimal resolution.

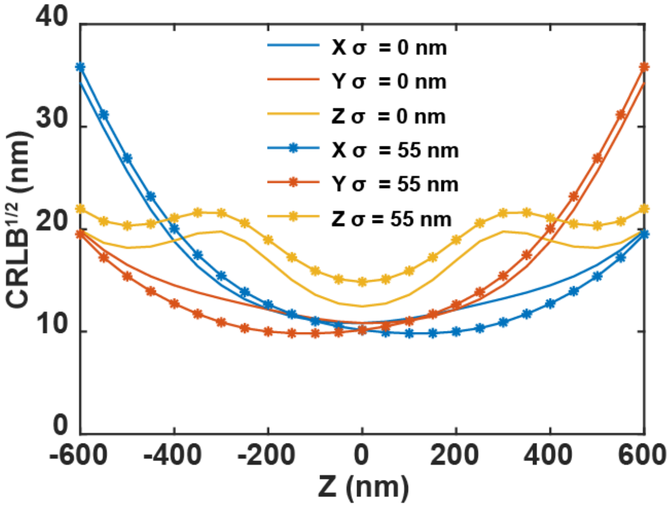

**SI Fig 1. Comparison of the impact of PSF Gaussian blurring on the theoretical CRLB.** The $\sigma$ corresponds to the $\sigma_x$ and $\sigma_y$ terms in SI Note 3 Eqs 3.8. When σ=0, Gaussian blurring is not applied. The pixel size of the data used is 330nm. The simulation parameters are the same as those in ED Fig. 1.

For the calculation of $CRLB_{3D}$, it is as follows[3],

$$CRLB_{3D} = \frac{1}{N_z}\sum_{i=1}^{N_z}\left(CRLB_{mean,x} + CRLB_{mean,y} + CRLB_{mean,z}\right) \quad (4.5)$$

Similarly, the calculation of $RMSE_{3D}$ is as follows,

$$RMSE_{3D} = \frac{1}{N_z}\sum_{i=1}^{N_z}\left(RMSE_x^2 + RMSE_y^2 + RMSE_z^2\right)^{\frac{1}{2}} \quad (4.6)$$

Where $N_z$ represents the number of z-positions within the axial range.

#### 5. Localization test

To evaluate the accuracy of the learned up sampled PSF model, we performed localization tests using the same bead data that were utilized during training. The accuracy was assessed by measuring the localization bias in the x, y, and z dimensions, with the localization bias in z is calculated from,

$$z_{bias,i,z} = z_{i,z} - z_{GT,i,z} \quad (5.1)$$

where $Z_{GT}$ represents the ground truth position, which corresponds to the stage position. Similarly, the localization bias in x/y is calculated from,

$$x_{bias,i,z} = x_{i,z} - \operatorname*{median}_{z} x_{i,z} \quad \text{and} \quad y_{\text{bias},i,z} = y_{i,z} - \operatorname*{median}_{z} y_{i,z} \quad (5.2)$$

where $x_{i,z}$ $and$ $y_{i,z}$ are estimated using MLE for the lateral localization of each 2D bead image, the subscripts $i$ and $z$ refer to the indices of the beads and the axial slices within each bead stack, respectively.

### References

1. Liu, S. *et al.* Universal inverse modeling of point spread functions for SMLM localization and microscope characterization. *Nat. Methods* **21**, 1082–1093 (2024).

2. Leutenegger, M., Rao, R., Leitgeb, R. A. & Lasser, T. Fast focus field calculations. *Opt. Express* **14**, 11277 (2006).

3. Fu, S. *et al.* Deformable mirror based optimal PSF engineering for 3D super-resolution imaging. *Opt. Lett.* **47**, 3031 (2022).

4. Li, Y. *et al.* Accurate 4Pi single-molecule localization using an experimental PSF model. *Opt. Lett.* **45**, 3765 (2020).

5. Babcock, H. P. & Zhuang, X. Analyzing Single Molecule Localization Microscopy Data Using Cubic Splines. *Sci. Rep.* **7**, 1–8 (2017).

6. Li, Y. *et al.* Real-time 3D single-molecule localization using experimental point spread functions. *Nat. Methods* **15**, 367–369 (2018).

7. Kay, S. M. *Fundamentals of statistical signal processing: estimation theory*. (Prentice-Hall, Inc, 1993).
